# Vision and locomotion shape the interactions between neuron types in mouse visual cortex

**DOI:** 10.1101/058396

**Authors:** Mario Dipoppa, Adam Ranson, Michael Krumin, Marius Pachitariu, Matteo Carandini, Kenneth D Harris

## Abstract

In the mouse primary visual cortex (V1), sensory responses are shaped by behavioral factors such as locomotion. These factors are thought to control a disinhibitory circuit, whereby interneurons expressing vasoactive intestinal peptide (*Vip*) inhibit those expressing somatostatin (*Sst*), disinhibiting pyramidal cells (Pyr). We measured the effect of locomotion on these neurons and on interneurons expressing parvalbumin (*Pvalb)* in layer 2/3 of mouse V1, and found inconsistencies with the disinhibitory model. In the presence of large stimuli, locomotion increased *Sst* cell responses without suppressing *Vip* cells. In the presence of small stimuli, locomotion increased *Vip* cell responses without suppressing *Sst* cells. A circuit model could reproduce each cell type’s activity from the measured activity of other cell types, but only if we allowed locomotion to increase feedforward synaptic weights while modulating recurrent weights. These results suggest that locomotion alters cortical function by changing effective synaptic connectivity, rather than only through disinhibition.

## Introduction

Neocortical interneurons can be divided into genetically distinguishable types, which are arranged in specific functional circuits (Jiang et al., 2015; Kepecs and Fishell, 2014; Markram et al., 2015; Pfeffer et al., 2013; Tasic et al., 2016; Tremblay et al., 2016; Zeisel et al., 2015). There is great interest in understanding how these circuits map to specific features of cortical operation, and how different interneuron types influence each other and ultimately shape the activity of excitatory neurons.

In recent years, a “disinhibitory model” has been proposed as a canonical feature of many cortical areas. This model rests on converging evidence that interneuron-selective interneurons, such as those expressing vasoactive intestinal peptide (*Vip*), principally target somatostatin (*Sst)* interneurons. This arrangement has been observed anatomically in the hippocampus (Acsady et al., 1996a; Acsady et al., 1996b), and functionally in auditory cortex (Pi et al., 2013), medial pre-frontal cortex (Pi et al., 2013), frontal association cortex (Garcia-Junco-Clemente et al., 2017) and visual cortex (Fu et al., 2014; Karnani et al., 2016a; Pfeffer et al., 2013). *Sst* neurons in turn inhibit most other cortical neuronal classes (with the exception of other *Sst* cells), including pyramidal (Pyr) cells (Jiang et al., 2015; Karnani et al., 2016b; Pfeffer et al., 2013). There is thus a potentially disinhibitory circuit involving *Vip* cells, *Sst* cells, and Pyr cells (Fu et al., 2014).

In sensory cortex, this disinhibitory circuit seems consistent with the modulation of sensory responses by behavioral state seen during arousal, whisking, and locomotion. In barrel cortex, for instance, whisking causes activity to increase in *Vip* cells, decrease in *Sst* cells, and increase in Pyr dendrites (Gentet et al., 2012; Lee et al., 2013). Furthermore, while the activity of Pyr cells correlates strongly with that of *Pvalb* cells, as predicted by the disinhibitory model it correlates weakly or negatively with the activity of *Sst* cells (Gentet et al., 2010; Gentet et al., 2012). In visual cortex, *Vip* cells are activated when animals run (Fu et al., 2014; Reimer et al., 2014), so a disinhibitory circuit might also explain the increases in visually-driven activity seen during locomotion (Ayaz et al., 2013; Erisken et al., 2014; Fu et al., 2014; Niell and Stryker, 2010).

The disinhibitory circuit, however, makes a key prediction that is unevenly supported by the data: that locomotion should decrease the activity of *Sst* neurons. One study confirmed the prediction, reporting that locomotion significantly (though mildly) reduced *Sst* neuron activity (Fu et al., 2014). Other studies, however, report the opposite effect or mixed effects (Pakan et al., 2016; Polack et al., 2013; Reimer et al., 2014).

Indirect evidence to support this prediction comes from visual cortex, where locomotion decreases the strength of a phenomenon, size tuning, that is thought to depend on *Sst* interneurons. Many V1 neurons exhibit size tuning: their responses are reduced when stimulus size increases beyond a preferred size. *Sst* interneurons are thought to contribute to size tuning because they integrate inputs from wide regions of cortex (Adesnik et al., 2012; Zhang et al., 2014) and because suppressing their activity reduces the strength of size tuning (Adesnik et al., 2012). If locomotion reduced *Sst* cell activity via inhibition from *Vip* cells, it should similarly reduce size tuning. This prediction is correct at least in the deep layers of V1 (Ayaz et al., 2013) and also in lateral geniculate nucleus (LGN, Erisken et al., 2014), perhaps because of cortical feedback. In both regions, locomotion decreases size tuning.

To test the predictions of the disinhibitory model, and more generally to understand the roles and relationships of different types of interneurons, we systematically investigated the effect of locomotion on visual cortical responses to stimuli of different sizes. We used two-photon microscopy to measure the activity of genetically-identified *Sst, Vip,* and *Pvalb* interneurons, together with pyramidal cells identified genetically or based on the skewness of their calcium traces.

Our results indicate that locomotor modulation of each cell class depends critically on the conditions of visual stimulation. Accordingly, subtle differences in experimental conditions can explain many of the apparent contradictions between previous studies. Our results, moreover, indicate multiple ways in which the effects of locomotion contradict the disinhibitory model. Modeling suggests that the complex interaction between locomotion, stimulus size, and cell class can be explained instead by a simple reweighting of feedforward vs. recurrent synapses.

## Results

We used two-photon imaging to measure the activity of Pyr neurons and of *Sst, Vip,* and *Pvalb* interneurons in mouse V1 (Figure 1 and Supplementary Figure 1). Mice were head-fixed and free to run on an air-suspended ball (Niell and Stryker, 2010) while viewing a grating in a circular window whose size ranged between 5° and 60° of visual angle (Figure 1A1). We selected for analysis only cells whose receptive field and orientation tuning matched the stimulus, and corrected for out-of-focus fluorescence (Supplementary Figure 2; Chen et al., 2013; Peron et al., 2015).

**Figure 1.**
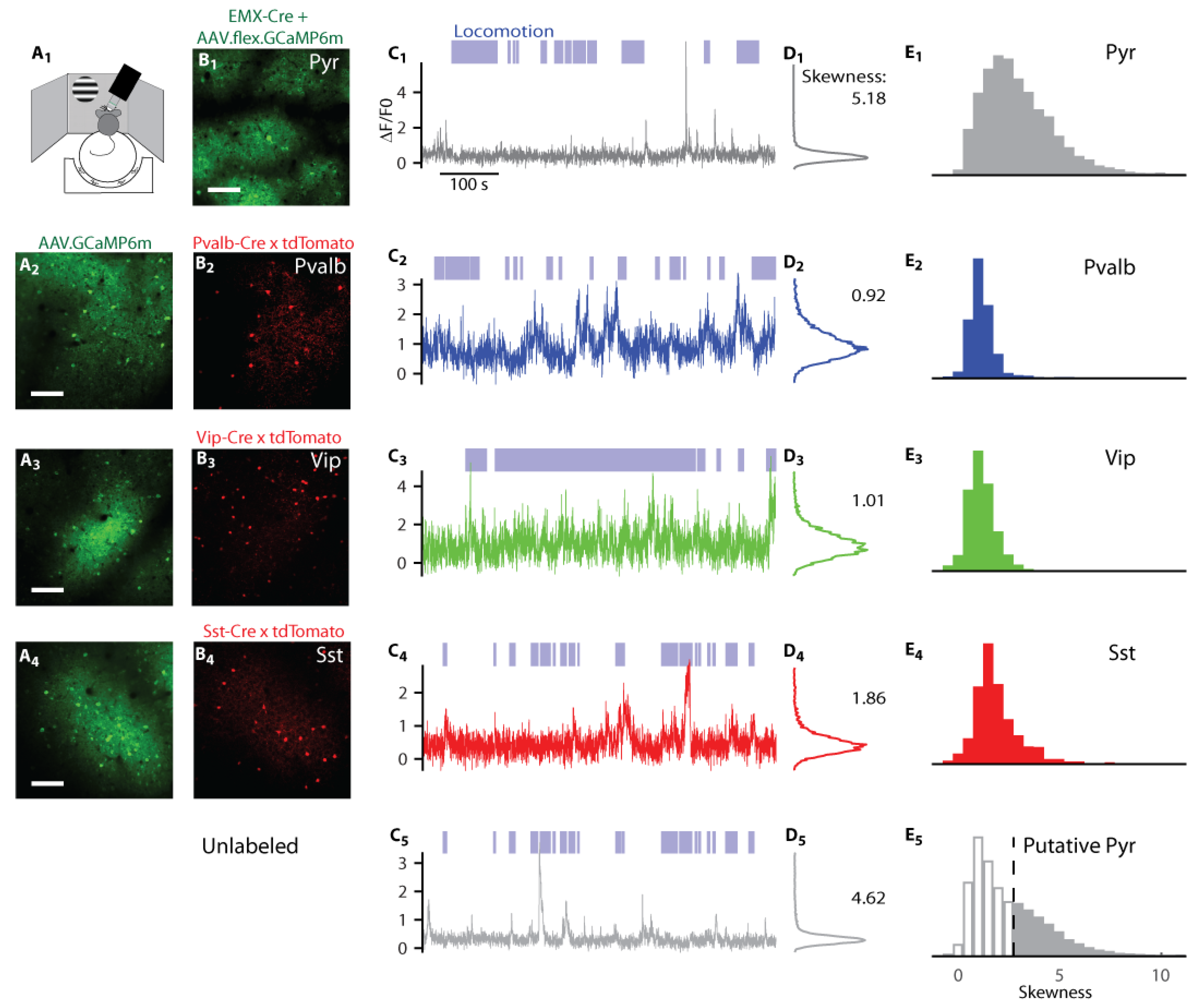
Recordings and classification of Pyr, *Pvalb, Vip*, and *Sst* cells in awake mouse V1.

A_1_) Experimental setup showing the air-suspended ball surrounded by the three screens for stimulus presentation.
A_2_-A_4_) Green fluorescence from three mice expressing GCaMP6 following virus injection. Scale bars: 100 μm.
B_1_) Green fluorescence from an *Emx1-Cre* mouse expressing GCaMP6m via virus injections.
B_2_-B_4_) Red fluorescence from the recordings in A_2_-A_4_, indicating tdTomato expression in *Pvalb* neurons (B_2_), in *Vip* neurons (B_3_), and in *Sst* neurons (B_4_).
C) Normalized fluorescent traces from five representative neurons of the Pyr, *Pvalb, Vip, Sst*, and unlabeled types. The unlabeled neuron in C_5_ was recorded simultaneously with the *Sst* example in C_4_. Blue shading above axes represents periods of locomotion (> 1 cm/s).
D) Histogram of fluorescence values for those five example neurons. The number indicates the skewness of the distribution.
E) Distribution of skewness values over all Pyr, *Pvalb, Vip, Sst*, and unlabeled neurons. Unlabeled cells above a skewness threshold of 2.7 (dashed vertical line) are classified as putative Pyr (E_5_).

### Recording the activity of identified cell classes

To identify neurons belonging to a specific class we used one of two approaches (Figure 1A, B). In the first approach (Figure 1A_2_-A_4_), GCaMP6m was virally expressed in all neurons, in mice where a specific cell class was also labeled with tdTomato. This approach allowed us to record the activity of identified interneurons and of many unlabeled neurons, which are likely to comprise mostly Pyr cells, along with interneurons of all classes except the labeled one. In the second approach, we expressed the calcium indicator exclusively in a chosen cell class, either by injecting a Cre-dependent GCaMP6m virus into an appropriate transgenic driver line (Figure 1B_1_ and Supplementary Figure 2B_2_) or via a triple-transgenic line that expressed GCaMP6s specifically in superficial layer Pyr cells.

**Figure 2.**
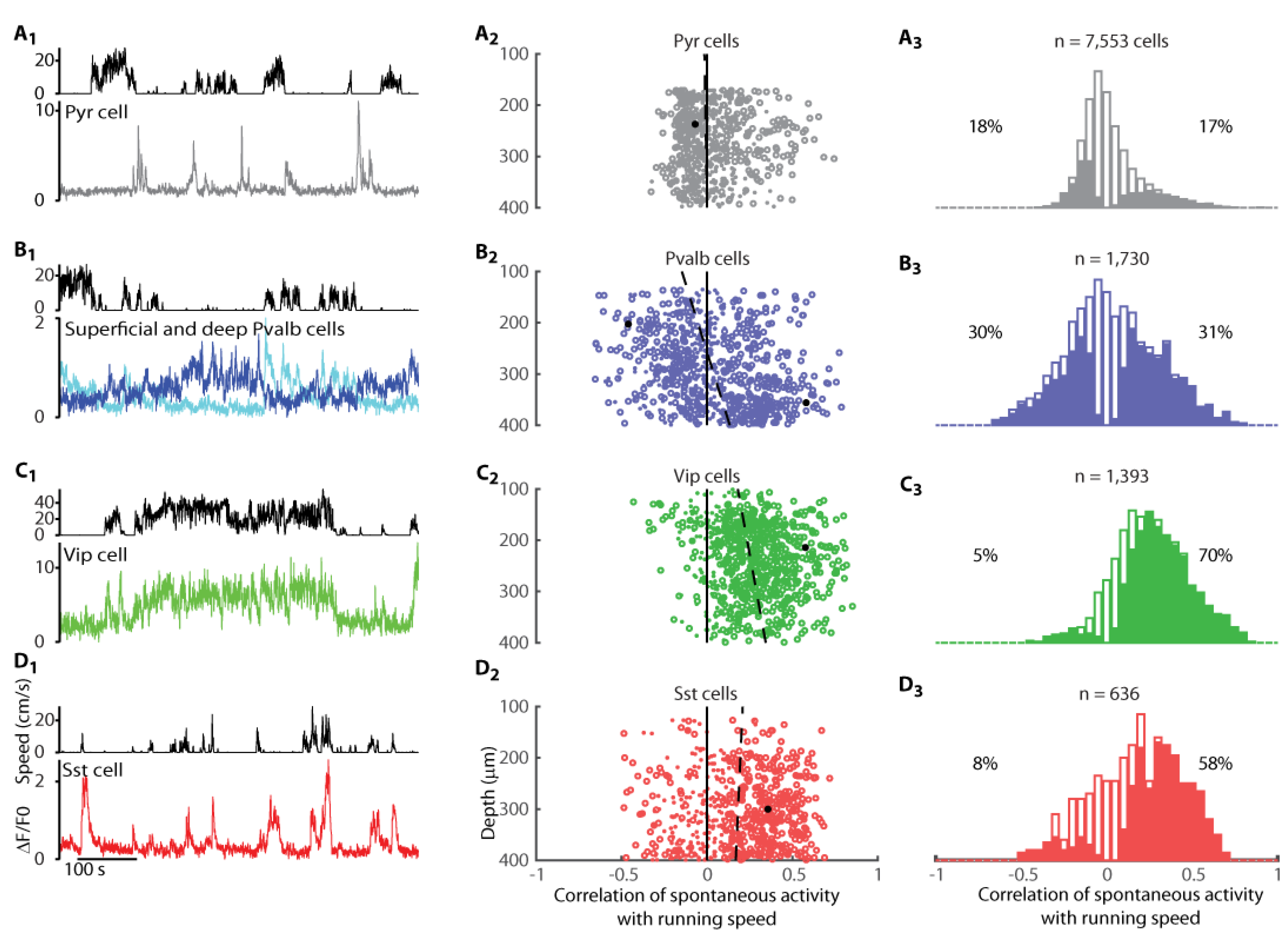
Correlates of running with neural activity in absence of visual stimuli (gray screen).

A_1_) Fluorescence of representative L2/3 pyramidal neuron (bottom) and simultaneous running speed trace (top).
A_2_) Correlation coefficient of recorded pyramidal cells with running speed, plotted vs. cell depth. Circles represent cells with significant correlations at p<0.05 (shuffle test); dots represent cells with insignificant correlations. For clearer visualization we plotted only a random subsample of 800 of genetically-identified Pyr cells. Dashed line represents fitted dependence of correlation vs. depth. Arrow indicates example cell shown in A_1_.
A_3_) Histogram of correlation coefficients of all pyramidal cells. Solid bars indicate significant correlations at p<0.05 (shuffle test). Values left and right of the histogram represent percentage of cells with a negative or positive correlation respectively.
B) Similar analysis for *Pvalb* neurons. The two traces in B_1_ (top) show fluorescence traces of representative *Pvalb* cells of upper and lower L2/3 (blue and cyan respectively).
C) Similar analysis for *Vip* cells.
D) Similar analysis for *Sst* cells.

Interneurons of all three classes fired much more frequently than pyramidal neurons (Figure 1C, D). Consistent with the sparse firing of superficial-layer pyramidal cells in mouse visual cortex (Niell and Stryker, 2010), the fluorescence traces of a typical identified pyramidal cell showed rare isolated calcium events (Figure 1C_1_). Because events with strong fluorescence were rare in these traces, the distribution of fluorescence was highly skewed (Figure 1D_1_). By contrast, the typical calcium fluorescence traces of identified *Pvalb, Vip,* and *Sst* interneurons showed frequent calcium events (Figure 1C_2_-C_4_), and the corresponding distributions of fluorescence showed little skewness (Figure 1D_2_-D_4_). Finally, many concurrently-measured unlabeled neurons showed sparse activity (example in Figure 1C_5_) and high skewness (Figure 1D_5_), resembling the identified Pyr cells (Figure 1C_1_, D_1_).

These differences in skewness were robust across the population, and allowed us to identify putative Pyr cells by their higher sparseness (Figure 1E). The precise value of the threshold for skewness made little difference to our results; we chose a conservative value of 2.7, that provided a small false-positive error rate (fraction of *Pvalb* neurons exceeding threshold: 24/1,511; *Vip:* 29/1,385; *Sst:* 91/537, Figure 1E_2_-E_4_), while still ensuring a majority of pyramidal cells exceeded threshold (2,598/4,949, Figure 1E_1_), for all methods of GCaMP expression (Supplementary Figure 3). Unlabeled neurons exceeding this threshold value are therefore likely to be pyramidal neurons, while cells below the value could be any cell type, and were therefore excluded from further analysis (Figure 1E_5_).

### Effects of locomotion on baseline activity

As a first test of the disinhibitory model, we asked how locomotion affected baseline activity, measured during long periods of gray screen presentation (Figure 2). In *Sst* cells, these measurements revealed ways in which previous apparently conflicting reports could be reconciled. In other interneurons, these measurements showed strong effects on baseline activity, an unexpected dependence on cortical depth, and phenomena that run opposite to the predictions of the disinhibitory model.

Consistent with previous results (Niell and Stryker, 2010; Polack et al., 2013; Saleem et al., 2013), the effects of locomotion on the baseline activity of identified Pyr cells were weak and diverse (Figure 2A). The sparse baseline activity (measured in response to a gray screen) of a typical *Pyr* cell changed only weakly with running speed (Figure 2A_1_). Across Pyr cells, the average correlation of baseline activity and running speed was close to zero (*ρ_gray_* = 0.03 ± 0.01 S.E.,(n = 7,553 identified Pyr cells, Figure 2A_3_). Nevertheless, 35% of Pyr cells showed a significant positive or negative correlation with speed (p < 0.05 shuffle test), significantly more than the 5% that would be expected by chance (p < 10^−16^, Fisher’s combined probability test). Similar results were seen in the putative Pyr neurons identified by their sparse firing (in unconditional labeling): correlations were small on average (*ρ_gray_* = -0.01 ± 0.01; SE, n = 5,666, Supplementary Figure 4A,B), but 49% of the cells were significantly positively or negatively correlated with speed (p < 0.05), significantly more than expected by chance (p < 10^−16^, Fisher’s combined probability test). In agreement with previous results obtained in darkness (Fu et al., 2014), when the monitors were switched off, the mean correlation of Pyr activity with speed was again close to zero (*ρ_dark_* = 0.00 ± 0.01, SE, Supplementary Figure 5A_1_,B_1_). However, cells tended to show similar effects of locomotion when monitors were turned on vs. off (Pearson correlation between *ρ_gray_* and *ρ_dark_* was 0.33; p < 10^−71^).

**Figure 3.**
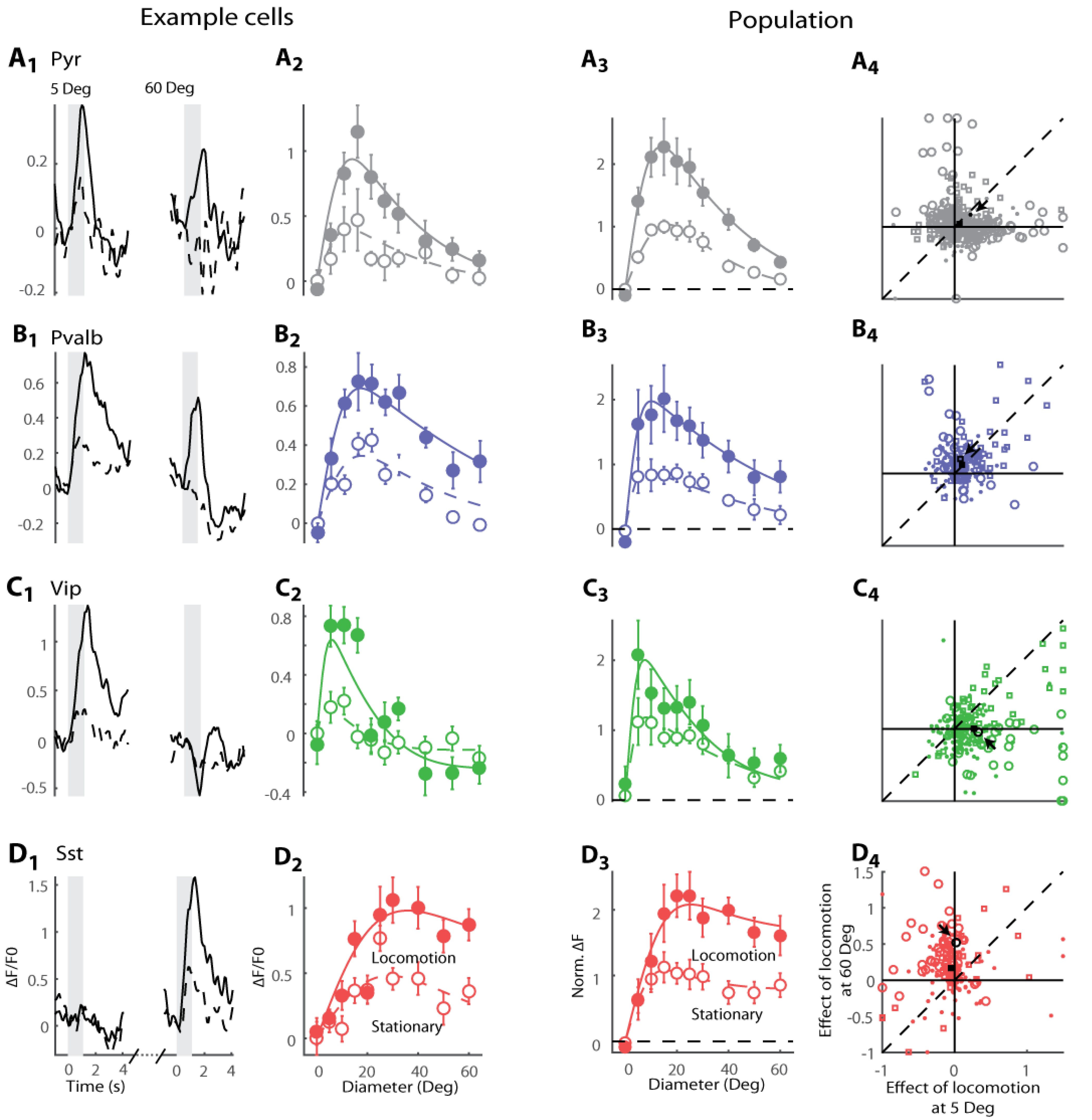
Locomotion increases visual responses for large drifting grating stimuli in *Sst* neurons, for small stimuli in *Vip* neurons, and for all stimulus sizes in *Pvalb* neurons.

A_1_) Responses of a representative Pyr neuron. Black curves show trial-averaged response in the stationary (dashed line) and locomotion (continuous line) conditions. Panels show responses to stimuli of 5° (left) and 60° (right). Gray shaded regions indicate the 1 s stimulus presentation period.
A_2_) Size tuning curve for this example cell. Solid line: locomotion; dashed line: stationary.
A_3_) Size tuning curve, averaged over Pyr cells after normalization. Solid line: locomotion; dashed line: stationary.
A_4_) Scatter plot showing change with locomotion of normalized responses to large stimuli (60°; y-axis) and to small stimuli (5°; xaxis). Circles represent cells whose responses have a significant interaction between size and locomotion (multi-way ANOVA over 5° and 60° stimuli only), squares represent cells that did not have a significant interaction but did have a significant effect of locomotion; dots represent cells with no significant effect of locomotion. Arrow marks example cell shown in A_1-2_, square marks mean response.
B-D) ,similar analysis for *Pvalb, Vip*, and *Sst* neurons.

**Figure 4.**
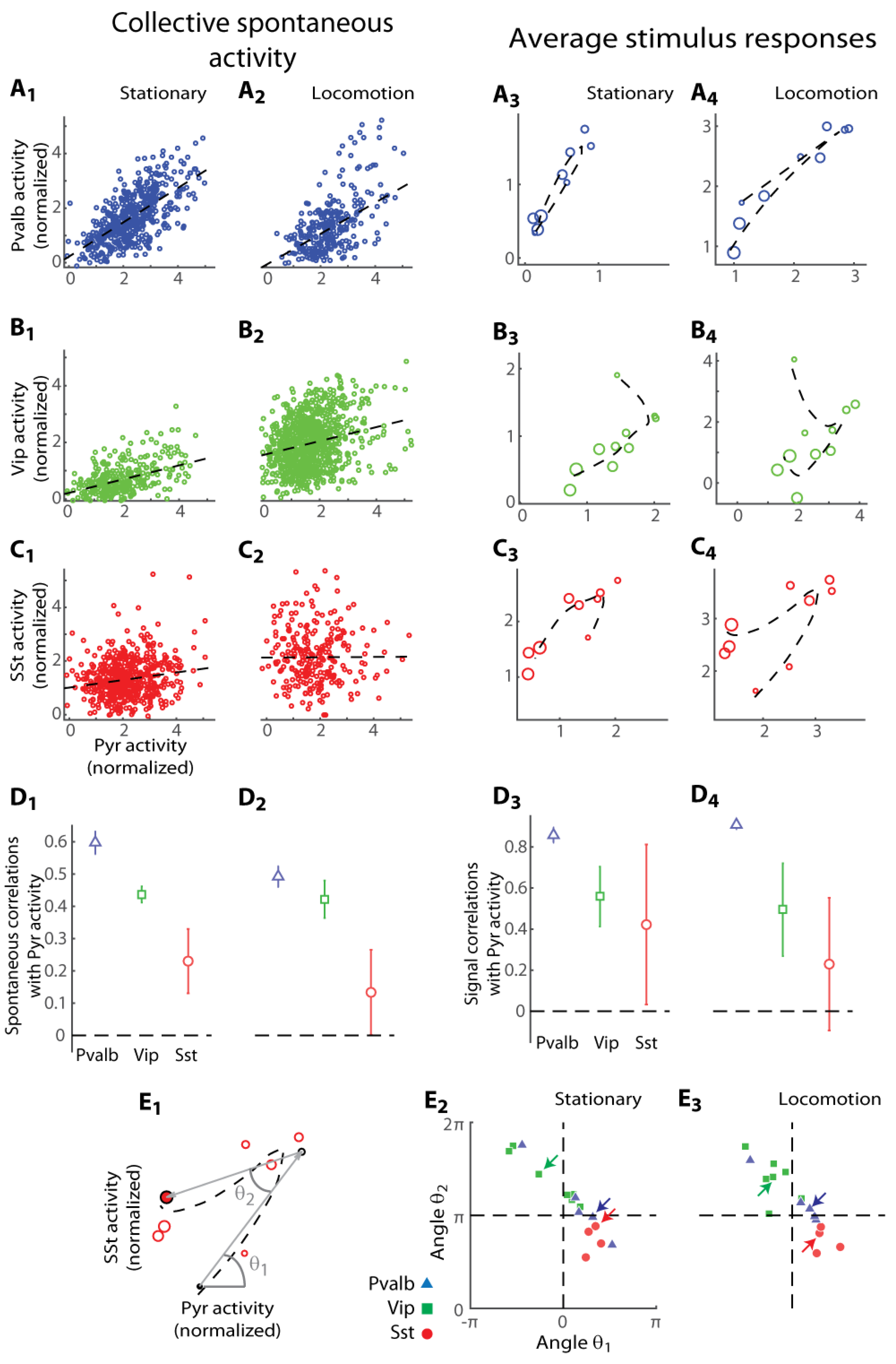
Pvalb cells are highly correlated with simultaneously recorded putative pyramidal cells, but Vip and Sst cells show nonlinear correlations.

A_1_ and A_2_) Summed activity of Pvalb population vs. Pyr population in the gray screen condition during stationary periods (A_1_) and during locomotion (A_2_). Each circle represents the simultaneous normalized value of the excitatory and the inhibitory populations at one time point. Dashed line indicates linear regression estimate of signal correlation.
A_3_ and A_4_) Average stimulus response of Pvalb population vs. average response of Pyr population in a typical experiment during stationary periods (A_3_) and during locomotion (A_4_). Each point represents a response to a stimulus, with larger circles representing larger stimuli. Dashed line represents non-linear interpolation of the Pvalb and Pyr size tuning obtained from their size-tuning curves.
B,C) Same as A, for Vip and Sst interneurons. Note the nonlinear signal correlations.
D_1_ and D_2_) Summary plots of spontaneous correlations during stationary periods (D_1_) and during locomotion (D_2_).
D_3_ and D_4_) Linear signal correlation (Pearson correlation coefficient) between the Pyr population and the three classes of interneurons averaged across all experiments during stationary periods (D_3_) and during locomotion (D_4_).
E_1_) Same plot represented in C_4_ illustrating the characteristic angles used to illustrate the nonlinear relationship between each interneuron class and Pyr mean visual responses. Circles with black contour indicate the minimum (5°, black filled) and maximum sizes (60°, red filled), and the cell’s preferred size (white filled). θ_1_ is the angle relative to horizontal of the line joining the response to 5° stimuli and the preferred stimulus for Pyr neurons. θ2 is the angle between the latter line and the line joining the response preferred size to the response to 60°.
E_2_ and E_3_) Angle θ2 vs. θ1 as defined in E_1_ for each experiment during stationary periods (E_2_) and during locomotion (E_3_). In E_3_ arrows indicate examples in (A_3_-C_3_); in E_4_ arrows indicate examples in (A_4_-C_4_).

**Figure 5.**
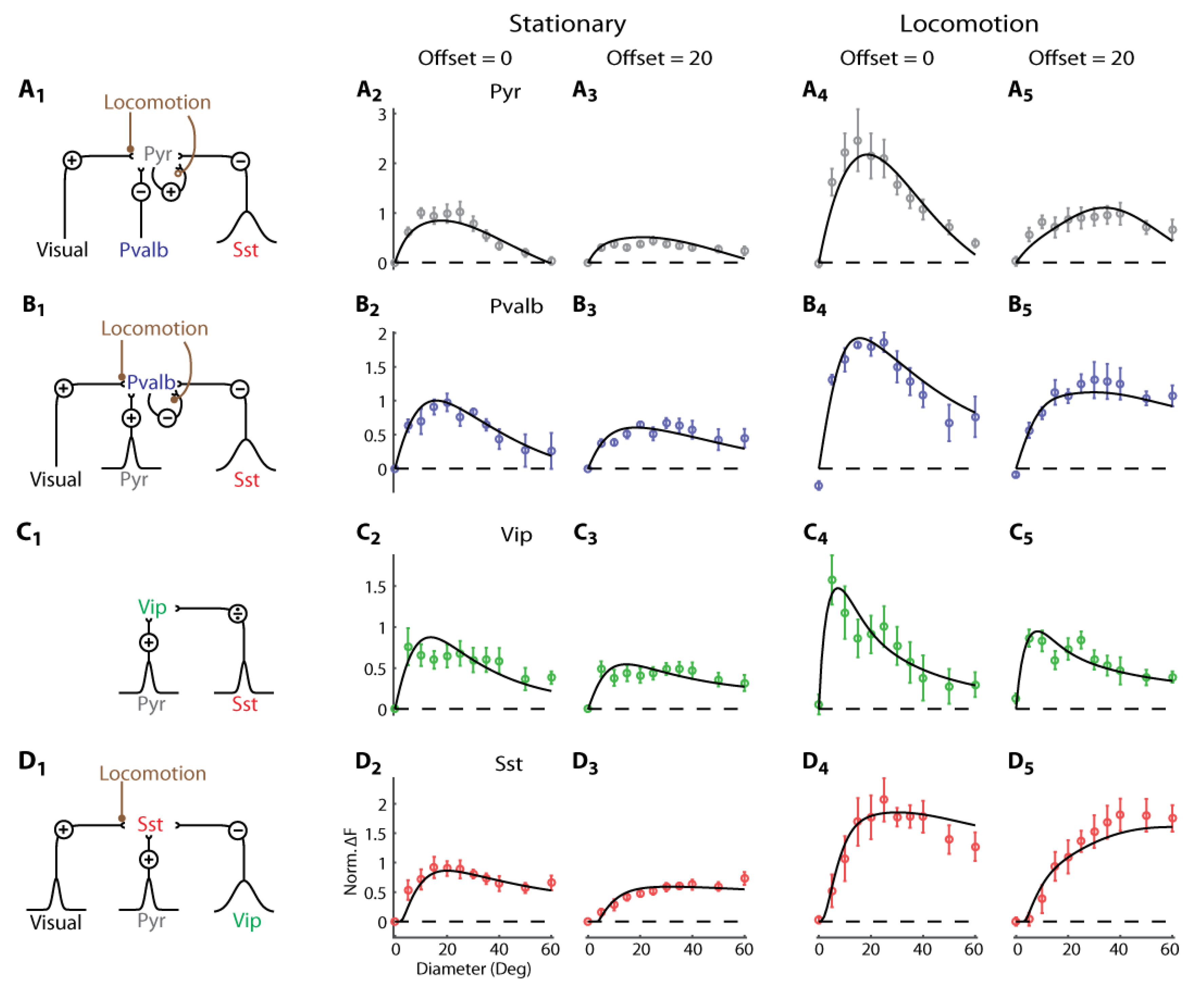
Computational model of size tuning in L2/3 of mouse V1. We searched the parameter space of neural field theories to find a model that can accurately reproduce our experimental results.

A_1_) Input synapses received by Pyr cells, and their modulation by locomotion. Pyr cells receive feedforward visual excitatory inputs (e.g. from thalamus or other cortical layers) and subtractive inhibition from *Pvalb* and *Sst* cells. Locomotion increases the synaptic weights from external visual inputs to Pyr cells (filled circles) while decreasing recurrent Pyr-to-Pyr connections (empty circles). Pyr cells integrate from a broad pool of *Sst* cells (large Gaussian curve).
A_2_-5) Model fit (black curves) of the size tuning data during stationary (A_2_,_3_) and locomotion (As) periods, visualized for centered cells (offset = 0, A_2_,4) and off-centered cells (offset = 20, A_3_,5).
B_1_) Input received by *Pvalb* cells. Locomotion boosts feedforward synaptic weights to Pvalb neurons while decreasing recurrent *Pvalb-to-Pvalb* synaptic weights.
B_2-5_) Same as (A_2-5_), for *Pvalb* cells.
C_1_) Input received by *Vip* cells. *Vip* cells receive divisive inhibition from *Sst* cells; no modulation of these synapses by locomotion is required to obtain a good fit.
C_2_-5) Same as (A_2_-5), for *Vip* cells.
D_1_) Input received by *Sst* cells. In addition to inputs from Pyr and *Vip* cells, *Sst* cells receive a feedforward input that we propose to originate from thalamus or other cortical layers. As with Pyr and *Pvalb* cells, the strengths of these synapses are boosted by locomotion.
D_2_-5) Same as (A_2_-5), for *Sst* cells.

The effects of locomotion on the baseline activity of *Pvalb* interneurons were stronger and more varied, and also depended on cortical depth (Figure 2B). For example, in two typical *Pvalb* cells imaged simultaneously (Figure 2B_1_), activity decreased with running speed in the more superficial cell (darker trace = -0.47, p < 0.01, shuffle test) and increased in the deeper cell (lighter trace, *ρ* = 0.58, p < 0.01 shuffle test). These results were typical of the population (n = 1,730), where the correlations were strong and depended significantly on depth (robust regression, p < 10^−21^; Figure 2B_2_), with high consistency across experiments (p < 0.018, t-test; Supplementary Figure 6). Among *Pvalb* cells in superficial L2/3 (depth <300 μm, n = 843) the average correlation with speed was slightly negative (*p_gray_* = -0.05 ± 0.03, SE), and this correlation was significantly negative in 36% of the cells and significantly positive only in 24% of the cells (p<0.05; shuffle test). The situation was reversed in deeper L2/3, where on average *Pvalb* cells showed a weak positive correlation with speed (*ρ_gray_* = 0.11 ± 0.04,SE, n = 831), with correlations significantly positive in 47% of cells and negative in only 18% of cells (p < 0.05; shuffle test). When pooling across depth, therefore, a wide variety of effects was seen (Figure 2B_3_), echoing the wide and bimodal range of correlations observed in darkness (Fu et al., 2014). When the monitors were switched off, the mean correlation of *Pvalb* activity with speed was slightly negative (*ρ_dark_* = -0.14 ± 0.01, SE, Supplementary Figure 5A_2_,B_2_) with a positive correlation between *ρ_gray_* and *ρ_dark_* across cells (Pearson correlation 0.49; p < 10^−22^).

Also as expected (Fu et al., 2014), the typical *Vip* cell increased baseline activity markedly with locomotion (Figure 2C_1_), and the overall population showed almost exclusively positive correlations with running speed, with a mean correlation of *ρ_gray_* = 0.27 ± 0.03 (SE, n = 1,393). The correlation increased significantly with cortical depth (robust regression; p < 10^−10^), an effect that was robust across experiments (p < 0.01, t-test; Supplementary Figure 6). We found similar results when we repeated these measurements with the monitors turned off (*ρ* = 0.30 ± 0.06, SE, Supplementary Figure 5A_3_,B_3_) with a positive correlation between *ρ_gray_* and *ρ_dark_* across cells (Pearson correlation 0.54; p < 10^−48^).

Contrary to the predictions of the disinhibitory circuit, however, locomotion generally increased the baseline activity of *Sst* interneurons (Figure 2D). The typical *Sst* cell increased its baseline activity markedly with locomotion (Figure 2D_1_) and across the population the correlation of baseline activity with running speed was on average positive (*ρ_gray_* = 0.18 ± 0.02, SE, n = 636, Figure 2D_3_), regardless of depth (robust regression, p = 0.39; Figure 2D_2_).

These effects of locomotion on the baseline activity of *Sst* interneurons confirm some previous results (Pakan et al., 2016; Polack et al., 2013) but they appear to disagree with other measurements (Fu et al., 2014). Indeed, these results run opposite to the predictions of the disinhibitory circuit, which would have predicted a negative effect of locomotion on baseline activity of *Sst* cells.

To confirm the validity of these results, we first ensured that they were not due to background fluorescence that might originate from other cell classes. We repeated the measurements in *Sst-IRES-Cre* mice where we expressed the calcium indicator only in *Sst* cells by injecting a Cre-dependent GCaMP6m virus (Supplementary Figure 2B_2_). At the time of the injection, the mice were already adult, thus excluding the off-target expression that might occur in cells expressing Cre only transiently during development (Hu et al., 2013). These experiments confirmed our results: the average correlation of baseline activity with running speed was positive (Supplementary Figure 2C-E) in all the locations where GCaMP6 was strongly expressed, be it cell bodies or neuropil. A pixel-by-pixel analysis further confirmed that these results did not depend on the method of ROI detection.

Previous studies have suggested that the modulation of spontaneous activity by locomotion can depend on the phase of the locomotion period, with stronger responses at locomotion onset (Vinck et al., 2015). However, for all cell types we found similar correlations between fluorescence and running speed after removing transition periods between locomotion and stationary periods from the analysis (Supplementary Figure 7).

We next asked if the disagreement on the effects of locomotion on the baseline activity of *Sst* cells could be due to different visual conditions used in different experiments (Pakan et al., 2016). The experiments of Fu et al. (2014) were performed in darkness, whereas in ours (Figure 2D) and those of Polack et al. (2013), the mouse faced a gray screen. We thus measured *Sst* baseline activity in darkness, and found that the effects of locomotion, albeit still diverse, were now overall negative (*ρ* = -0.07 ± 0.02, SE; across experiments: p = 0.019, t-test, Supplementary Figure 5A_4_,B_4_,C). The same cell could show different modulation by locomotion depending on screen illumination (Supplementary Figure 5B4): for example, of the cells showing significant modulation in both conditions, 26% showed *ρ_dark_* < 0 and *ρ_gray_* > 0. Not all cells, however, showed this diversity. On average, in fact, *Sst* cells showed a positive correlation between *ρ_gray_* and *ρ_dark_* (Pearson correlation 0.34; p < 10^−8^).

These measurements, therefore, reconcile the apparent divergence of previous results (Fu et al., 2014; Pakan et al., 2016; Polack et al., 2013): the effect of locomotion on baseline activity of *Sst* cells is overall positive when mice view a gray screen, and mildly negative when mice are in darkness. The observations made with the gray screen, however, contradict the disinhibitory model.

### Effects of locomotion on responses to gratings of different sizes

Having witnessed the importance of controlling visual stimulation when studying the effects of locomotion, we next asked how these effects depend on stimulus size. We measured responses to drifting gratings (relative to baseline activity measured before stimulus onset), and we focused on cells that were visually responsive (significant effect of stimulus size, p < 0.05, 1-way ANOVA) and whose RFs were centered within 10° of the stimulus center. To discount possible effects of eye movements (whose occurrence might change during locomotion), we only selected trials where the pupil pointed within 10° from the average position. Similar results were found when we removed this restriction (Supplementary Figure 8).

Identified L2/3 Pyr cells were selective for small stimuli (in agreement with Adesnik et al., 2012), and exhibited, on average, a mild effect of locomotion (Figure 3A). A typical Pyr neuron responded substantially more to a 5° stimulus than to a 60° stimulus, regardless of locomotion (Figure 3A_1_). Accordingly, the neuron showed clear selectivity for smaller stimuli (Figure 3A_2_), similar to that seen in the overall population of identified Pyr cells (n = 1,250; Figure 3A_3_) where cells preferring large stimuli were rare (Supplementary Figure 9A). On average, locomotion slightly increased responses to both small stimuli (p < 10^−16^, paired t-test across cells, p < 0.01, paired t-test across experiments) and large stimuli (p < 10^−10^ and p = 0.02). However, this effect was diverse among cells, with locomotion significantly increasing or decreasing responses in 17% and 3% of Pyr cells respectively (p < 0.05; 2-way ANOVA, main effect of locomotion over 5° and 60° stimuli, Figure 3A_4_). Many cells (17%) showed a significant interaction of locomotion and stimulus size (p < 0.05; 2-way ANOVA over 5° and 60° stimuli; Figure 3A_4_; Supplementary Figure 10A,B_1_). In these cells, locomotion changed the relative response to large and small stimuli, as seen previously in deeper layers (Ayaz et al., 2013). Similar results were found in putative pyramidal cells identified by the sparseness of their calcium traces (Supplementary Figure 4C,D).

*Pvalb* interneurons were similarly selective for stimulus size, but showed a stronger and overwhelmingly positive effect of locomotion (Figure 3B). A typical *Pvalb* interneuron responded strongly to small stimuli and more weakly to larger stimuli, and its responses markedly increased during locomotion (Figure 3B_1_-B_2_). These effects were highly consistent across *Pvalb* interneurons (n = 277, Figure 3B_3_), with locomotion increasing firing rate in practically all cells (Figure 3B_4_). This effect was seen in responses to both large stimuli (p < 10^−n^, paired t-test across cells, p < 0.01, paired t-test across experiments) and small stimuli (p < 10^−13^ and p = Example cells 0.02), with no significant interaction between stimulus size and locomotion (p = 0.23, 2-way ANOVA over 5° and 60° stimuli; Supplementary Figure 10A,B_2_).

The responses of *Vip* interneurons (n = 239) were selective for stimulus size and increased with locomotion, but in contradiction with the disinhibitory circuit, this increase was generally restricted to responses to small stimuli (Figure 3C). A typical *Vip* interneuron responded most strongly to small stimuli during locomotion (Figure 3C_1_-C_2_). Similar results were seen across the population: *Vip* interneurons showed clear size tuning, and locomotion increased their responses to 5° stimuli (p < 10^−15^, paired t-test across cells, p = 0.04, paired t-test across experiments) but not to 60° stimuli (p = 0.82 and p = 0.86, Figure 3C_3_) with a significant interaction of size and locomotion (p < 10^−10^, 2-way ANOVA over 5° and 60° stimuli; Figure 3C_4_; Supplementary Figure 10A,B_3_). Similarly to baseline activity, the effects of locomotion on visual responses grew with cortical depth (robust regression, p < 10^−3^).

*Sst* interneurons tended to prefer large stimuli, if not the largest ones, and their responses – contradicting the disinhibitory circuit – increased with locomotion (Figure 3D). As observed by Adesnik et al. (2012), a typical *Sst* interneuron responded best to large stimuli, and even more so while the animal was running (Figure 3D_1_, D_2_). Similar results were seen across the population (n = 191, Figure 3D_3_), with overall activity peaking for ~15° stimuli during stationary conditions, and ~25° stimuli during locomotion. *Sst* cells showed a significant interaction between stimulus size and locomotion (p < 10^−7^, 2-way ANOVA over 5° and 60° stimuli), consistently across experiments (p < 0.01, t-test, Supplementary Figure 10A4) and mice (p < 0.01, t-test, Supplementary Figure 10B4). While locomotion did not significantly affect the responses to small stimuli (p = 0.06, paired t-test across cells, p = 0.65, paired t-test across experiments, Figure 3D_4_), it strongly increased the responses to large stimuli (p < 10^−9^ and p = 0.02), thus contradicting the disinhibitory circuit.

Some *Sst* cells, however, did show size tuning (Figure 3D, Supplementary Figure 9C,D). This observation is consistent with observations in anesthetized mice (Pecka et al., 2014), but differs from those of Adesnik et al. (2012) in awake mice. We reasoned that it may reflect high sensitivity of these cells to stimulus centering, and to investigate this possibility, we studied how size tuning varies with the distance between receptive field center and stimulus center. Regardless of cell class, when the stimulus was distant, the cells preferred the largest stimulus (Supplementary Figure 11). Size tuning emerged when stimulus distance decreased. For *Sst* cells, in particular, size tuning appeared when stimuli were < 20° away from the receptive field center (Supplementary Figure 11D).

For all cell types, we saw similar interactions of size tuning and locomotion when cells were recorded in the binocular and monocular regions of visual cortex (Supplementary Figure 12). Furthermore, the results did not depend on whether or not we deconvolved the calcium traces to estimate spike rates (Supplementary Figure 13).

In summary, our results for large stimuli contradict the disinhibitory model, which predicts that locomotion should increase firing in *Vip* cells and decrease it in *Sst* cells. Instead, we saw something closer to the opposite behavior: in the presence of large stimuli, locomotion does little to *Vip* cells and boosts the activity of *Sst* cells.

### Correlations of interneuron activity with putative pyramidal activity

To further test the disinhibitory model, we examined the patterns of correlation between interneuron types and Pyr cells (Figure 4). To measure correlation, we relied on our ability to record simultaneously the activity of identified interneurons and of putative Pyr neurons (unlabeled sparse-firing cells, with skewness > 2.7, Figure 1). This analysis revealed dramatic differences between interneuron classes.

Consistent with the view that *Pvalb* interneurons track the activity of Pyr cells (Cruikshank et al., 2007; Isaacson and Scanziani, 2011; Okun and Lampl, 2008; Ozeki et al., 2009; Renart et al., 2010), we found strong positive correlations between *Pvalb* and putative Pyr populations (Figure 4A). In each experiment, we summed the activity of the *Pvalb* interneurons and compared it to the summed activity of the simultaneously recorded putative Pyr cells. Only cells whose receptive field lay within 10° from the stimulus center were considered. In a typical experiment the spontaneous correlations (*ρ*_0_) measured in the presence of a gray screen were strongly positive, whether the mouse was stationary (*ρ*_0_ = 0.70, Figure 4A_1_) or running (*p*_0_ = 0.60, Figure 4A_2_). Similar results were seen across experiments (*ρ*_0_ = 0.60 ± 0.04, SE, during rest and 0.49 ± 0.03 during locomotion). Examining population signal correlations – i.e. the relationship of the mean summed responses of the *Pvalb* and Pyr cells across stimuli – also showed a strong, positive correlation in both stationary (*ρ_s_* = 0.95 for the example in Figure 4A_3_, 0.86 ± 0.04 across all experiments) and locomotion conditions (*ρ_s_* = 0.97 for the example in Figure 4A_4_, 0.91 ± 0.02 SE across all experiments). Similar results were found for population noise correlations (i.e. the relationship between trial-to-trial variability in summed activity of the *Pvalb* and Pyr populations; Supplementary Figure 14). These results were not biased by the exclusion of low-skewed Pyr cells, which did not pass our conservative skewness threshold: when we lowered the threshold (therefore including more Pyr cells but also some unlabeled inhibitory neurons), the correlations of putative Pyr cells with *Pvalb* cells remained high (Supplementary Figure 15A,B,C).

*Vip* cells showed a markedly different behavior (Figure 4B). Their correlations with putative Pyr populations differed from those of *Pvalb* cells in two respects. First, while spontaneous and noise correlations tended to be positive (*ρ*_0_ = 0.43 ± 0.03, SE, during stationarity and 0.42 ± 0.06 during locomotion; *ρ_n_* = 0.33 ± 0.07, SE, during stationarity and 0.37 ± 0.02 during locomotion), they were weaker than those of *Pvalb* cells, at least during stationarity (Figure 4D1 and Supplementary Figure 14). Second, the relationship between the population mean responses of *Vip* and putative Pyr cells was clearly nonlinear (Figure 4B3,4). This nonlinearity reflects the different size tuning of *Vip* and Pyr cells, with *Vip* responses peaking at smaller stimulus sizes than Pyr responses (compare Figure 3C_3_ and A_3_). It further suggests that, unlike for *Pvalb* cells, the sensory tuning of *Vip* cells during locomotion cannot be explained by a simple tracking of excitatory activity.

*Sst* interneurons showed yet a different sort of behavior, which depended strongly on locomotion (Figure 4C). The correlation of the *Sst* and Pyr populations was positive in stationary conditions (*p*_0_ = 0.25 for the example in Figure 4C1, 0.23 ± 0.10 SE across experiments) but weak during locomotion (*p*_0_ = -0.01 for the example in Figure 4C2, 0.13 ± 0.13 SE across experiments). Noise correlations were also positive (Supplementary Figure 14; *p_n_* = 0.39 ± 0.11 SE, stationary, *ρ_n_* = 0.30 ±0.04 SE, locomotion). Signal correlations showed a non-linear character, but this differed to that of *Vip* cells (Figure 4C_3_,_4_).

To quantify the form of nonlinear relationship between each interneuron class’s mean responses and those of putative Pyr cells, we parameterized their relationship using two angles θ1 and θ2 Figure 4E_1_). These angles were similar in stationary and locomotion conditions, and were consistent across datasets (Figure 4E); they were also robust to changes in the skewness threshold (Supplementary Figure 15D,E). The different form of nonlinear relationship between *Sst* and *Vip* cells result in a significant difference in 01 (stationary: p = 0.026, locomotion: p < 0.01, Watson’s U^2^ permutation test, N=1,000 permutations). Similarly, there was a significant difference of 02 between *Sst* and *Vip* cells (stationary: p = 0.039, locomotion: p = 0.036, Watsons U^2^ permutation test, N=1,000 permutations).

Overall, thus, these results paint a more complex picture than would be expected from the disinhibitory model. The model predicts that Pyr activity should increase with *Vip* activity and decrease with *Sst* activity. This prediction is consistent with correlations measured during spontaneous activity in darkness, but not with the covariation of stimulus responses seen in the presence of medium to large stimuli, where both *Vip* and *Sst* cells decrease their activity with stimulus size. Visual stimulation, therefore, creates situations where the disinhibitory circuit cannot account for size tuning.

### Recurrent network model

To explore how size tuning of the different neuron classes in layer 2/3 of area V1 could arise from cortical circuitry, we asked if their size tuning curves could be reproduced by a recurrent network model. We fit the model with a novel approach: we estimated the synaptic input parameters for each class of neurons to be those optimally predicting that class’ average population sensory responses, with the average population activity of all other cell classes clamped to their measured values.

We modeled the activity of each cell class by a “neural field”: a number that varied across the retinotopic cortical surface, representing the mean firing rate of all cells of that class at that location. The sensory response of cell class *α* was modeled as a function 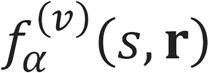 of stimulus size *s* and retinotopic position r (relative to stimulus center); dependence on r was assumed to be circularly symmetric. The superscript *v* indicates the locomotion condition (*v* = 0: rest, *v* = 1: locomotion). Synaptic connections were modeled as functions of retinotopic location: the connection strength to a cell of type *α*_1_ at location ***r***_1_ from a cell of type *α*_2_ at location *r*_2_ decays as a two-dimensional Gaussian function of the distances between receptive field centers 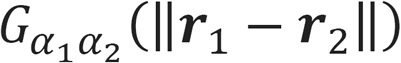. The strength and spatial spread of these connections were fit by exhaustive search, minimizing the squared error between the predicted and actual rates summed over all stimulus sizes and offsets and locomotion conditions. Each cell integrates its inputs using a threshold-linear activation function. For each inhibitory connection, we tested both divisive and subtractive inhibition, with the choice made automatically for each inhibitory synapse type to minimize errors. The firing of dLGN inputs *h(s, r)* was modeled using a ratio of Gaussians (Ayaz et al., 2013), with parameters (for both rest and locomotion conditions) fit to the data of Erisken et al. (2014).

The model accurately predicted the size-tuning curves of each cell class, for both centered and off-centered stimuli (Figure 5), but this success was predicated on certain conditions. Specifically, to predict the strong response of *Sst* cells to large, off-center stimuli (Figure 5D_3_ and D_5_), the model needed to include external excitatory input to these cells (e.g. from thalamus or other from cortical layers), because large stimuli elicited little response in Pyr cells (Figure 5A_2_-A_5_). A good fit was only obtained if this feedforward input to *Sst* cells had broad size tuning, as would be seen in thalamic neurons or perhaps in excitatory neurons of other cortical layers (Figure 5D_1_). Moreover, to obtain the observed similarity in tuning of *Pvalb* and Pyr cells (Figure 5A,B), the model required these cell classes to have similar inhibition from *Sst* cells. Furthermore the model required Pyr neurons to lack *Vip* input. Finally, our parameter search only gave good results with divisive inhibition from *Sst* to *Vip* cells (Figure 5C_1_): subtractive inhibition could not produce the observed sharp size-tuning favoring small stimuli (Figure 5C_2_-C_5_).

We next asked what modifications of the model parameters could explain the effects of locomotion on the sensory responses of each cell type. Modeling the locomotor modulation of size tuning required that locomotion change synaptic strengths: if we kept the input synapses of Pyr cells at the strengths fit to stationary data, while clamping the activity of other cell types to their values during rest, we obtained a poor prediction of Pyr cells’ size tuning during locomotion (not shown). To capture the effects of locomotion on each cell type, we therefore searched for all possible ways that locomotion could modulate the thalamocortical and recurrent synaptic weights of each class (Supplementary Figure 16).

For Pyr cells, we could prove analytically that the fit quality depended only on the strength of their “effective connections” (see Experimental Procedures). These effective connections depended on several network parameters, including the strengths of external, recurrent excitatory, *Pvalb,* and *Sst* inputs onto Pyr neurons, and take into account the amplification caused by recurrent excitation and *Pvalb* inhibition (Douglas et al., 1995). Thus, while the model fit identified unique values of the effective connections, these values could in turn be achieved through multiple possible strengths of the synaptic parameters (Prinz et al., 2004).

Capturing the modulation of Pyr tuning by locomotion required an increase in effective connection from external inputs to Pyr cells, and a decrease in effective connection of *Sst to* Pyr cells. Together, this produced the observed strong increase in responses of centered cells to medium-sized stimuli, together with a milder increase in response to larger stimuli (Figure 5A_2_ vs. A4). However, this change in effective connection strength did not require a weakening of the physical Sst–>Pyr connections: the same change in effective connection could also be achieved by an increase in external excitation together with a decrease in recurrent excitation (Supplementary Figure 16A2). Intriguingly, these two effects are precisely those observed *in vitro* during cholinergic modulation of cortical synapses (Gil et al., 1997).

Consistent with the close correlations of *Pvalb* and Pyr cells, modeling the observed effects of locomotion on *Pvalb* size tuning required similar modulations to those required for Pyr cells: a decrease in effective inhibition from *Sst* cells. Again, however, this did not necessarily require a weakening of *Sst–>Pvalb* synapses, as the same effect could be obtained via strengthening of *Pvalb->Pvalb* connections and of the external input. Producing the observed effects of locomotion on *Vip* cell tuning required no further changes in effective connection: locomotion only increased the responses of centered *Vip* cells to small stimuli (Figure 5C_2_ vs. C_4_), and this increase could be readily provided by increased activity of local Pyr cells. Finally, correct modulation of *Sst* firing required boosting of the external excitatory inputs responsible for their responses to large stimuli (Figure 5D), but did not require any change to their inhibitory inputs.

In summary, we were able to capture the effects of locomotion on all cell types through a reweighting of feedforward and recurrent connections: an increase in external excitatory input to all cell types, a decrease in recurrent excitation between Pyr cells, and an increase in recurrent inhibition between *Pvalb* cells.

## Discussion

We have shown that locomotion does not simply increase or decrease the activity of a particular cell class: its effects depend on the precise sensory conditions, and even on cortical depth. Its effect on sensory responses was an increase in all cell classes, but the increase varied with cell type and stimulus, being largest in *Sst* responses to large stimuli and in *Vip* responses to small stimuli. The effects of locomotion on baseline neural activity (as assessed by gray screen viewing) were more complex: locomotion increased activity in most *Sst* and *Vip* neurons, and had diverse effects on *Pvalb* and Pyr cells, suppressing most *Pvalb* cells in superficial L2/3 and increasing activity in deeper *Pvalb* cells.

Our results further indicate that apparent discrepancies in the literature may in fact reflect the subtly different methods used in different experiments. Specifically, two studies on the spontaneous activity of *Sst* cells reported opposite effects of locomotion: increased activity (Polack et al., 2013) and decreased activity (Fu et al., 2014). When we replicated their experimental conditions (gray screen for the first study, complete darkness for the second), we replicated both observations in a common set of neurons. These results reconcile the apparent contradiction between these studies (see also Pakan et al., 2016).

The results also reinforce the importance of correcting for out-of-focus fluorescence in 2-photon calcium imaging. Indeed, without correcting for this confound, one would observe an artefactual negative correlation of fluorescence with running speed, particularly in image regions with weak GCaMP expression (Supplementary Figure 2).

We had set out to test a specific, simple “disinhibitory” hypothesis: that during locomotion, increased *Vip* activity would inhibit *Sst* cells, and thus increase the activity of pyramidal cells (Fu et al., 2014). Some of our results were consistent with the hypothesis. For instance, we confirmed that locomotion increases *Vip* responses. Other results, instead, could not be explained by disinhibition alone. The hypothesis predicts that increased *Vip* activity should suppress *Sst* firing, and yet we found that *Sst* firing increased during locomotion, as long as visual inputs were present (even just a blank gray screen).

Another area where previous models need to be updated concerns the mechanisms of size tuning. Adesnik et al (2012) proposed that an important contribution to size tuning comes from the activity of *Sst* cells. They reported these cells as firing mostly during locomotion and showing negligible size tuning. Our experiments confirmed that *Sst* cells fire more during locomotion, but replicated the finding about size tuning only when stimuli were poorly centered on the receptive field. When they were centered, instead, *Sst* neurons displayed marked preferences for intermediate stimulus sizes. In light of this finding, the role of *Sst* neurons in size tuning needs to be re-evaluated. Size tuning is likely to reflect a more complex interaction between the different cell classes.

To understand what form this interaction might take, we built a network model and searched for the parameters that caused it to produce accurate fits to our measurements. Previous studies that proposed a network circuit to explain surround suppression in V1 had not included different cell types (Ozeki et al., 2009; Rubin et al., 2015) or had not validated the predictions with quantitative fits (Litwin-Kumar et al., 2016). Our analysis cast further doubt on the possibility that a disinhibitory mechanism is sufficient to explain changes in size tuning. Indeed, had a disinhibitory mechanism been sufficient, the measured locomotion modulation of interneuron tuning – with no changes in synaptic connection strengths – would have been sufficient to cause the observed modulation in pyramidal cell tuning. Instead, the model required locomotion to change effective connections between neuronal classes, i.e. the effect of one class on another after taking into account amplification through recurrent excitation and inhibition.

These specific changes in effective connections could in turn be instantiated through multiple possible modulations of physical synaptic strengths. The activity produced by a neural circuit is not always sufficient to constrain its underlying connectivity (Prinz et al., 2004); for the current model, we could mathematically prove that multiple underlying connectivity patterns yield identical sensory responses. Nevertheless, the parameter space consistent with our experimental observations favored one particularly attractive possibility, where locomotion would increase external excitatory input to all cell types, decrease recurrent excitation between Pyr cells, and increase recurrent inhibition between *Pvalb* cells. The first two of these are known effects of cholinergic modulation on cortical circuits (Gil et al., 1997).

The model required specific conditions to produce an accurate quantitative fit to our measurements, which we therefore consider experimental predictions. First, the model required that *Sst* cells receive a feedforward sensory input: i.e. an excitatory input conveying visual input other than from local Pyr cells. This was required because off-center *Sst* cells fire strongly to large stimuli, but neither centered nor off-center Pyr cells fire strongly enough to large stimuli to provide sufficient input. Whether *Sst* cells receive direct thalamic inputs is controversial (Cruikshank et al., 2010; Lee et al., 2013; Tan et al., 2008). Even without direct thalamic afferents however, such an input could be conveyed to superficial *Sst* cells via other cortical layers. Interestingly, the optimal model parameters required that the external inputs *Sst* cells receive be spatially diffuse, as would be expected if this input had experienced an additional round of divergence through Pyr cells of other layers. A second condition required by our model was that *Vip* cells receive divisive, rather than subtractive inhibition from *Sst* cells. We are not aware of any data on this question, and therefore consider it to be a model prediction.

In contrast to the complex relationship of *Vip* and *Sst* neurons to summed Pyr activity, the activity of *Pvalb* interneurons was in all cases tightly locked to the local excitatory population. These data are therefore consistent with a primary role for *Pvalb* cells of stabilize the activity of the local circuit via tracking summed excitatory firing (Cruikshank et al., 2007; Ozeki et al., 2009; Renart et al., 2010), rather than directly sculpting receptive field shapes. In our model, we were able to reproduce the similar tuning of *Pvalb* and Pyr cells only if they received similar inputs from other inhibitory classes. Consistent with *in vitro* observations (Pfeffer et al., 2013), the model required that both Pyr and *Pvalb* cells received inhibition from *Sst,* but not *Vip* cells.

What computational benefit might the visual cortex derive from these interactions of locomotion, stimulus size, and cell type? One can consider two hypotheses for why locomotion should modulate visual cortical activity. In the first hypothesis, boosting of activity in visual cortex in running animals serves as a form of modality-specific attention: because vision is particularly important during navigation, locomotion increases V1 activity while suppressing the activity of competing sensory systems. This possibility is supported by the fact that locomotion appears to decrease activity in other cortical regions (Schneider et al., 2014; Zhou et al., 2014), but it cannot explain why locomotion has such diverse effects on V1 pyramidal cells, boosting some while suppressing others, and increasing responses only to certain sensory stimuli (Keller et al., 2012; Saleem et al., 2013). The second hypothesis holds that locomotor modulation of V1 is not simply a matter of gain control, but a complex and diverse neuronal integration of visual and locomotor input. Indeed, work in virtual reality environments has suggested that, rather than simply making visual responses larger or smaller, locomotor modulation may underlie more complex computations, such as integrating the animal’s own movement with that of the world (Keller et al., 2012; Saleem et al., 2013). Diverse interaction of sensory stimuli and locomotion in different interneuron classes might form a key mechanism behind this integration.

## Experimental procedures

All experimental procedures were conducted according to the UK Animals Scientific Procedures Act (1986). Experiments were performed at University College London under personal and project licenses released by the Home Office following appropriate ethics review.

### Mice

Experiments in which an interneuron class was labelled with tdTomato and recorded together with other cells were conducted in double-transgenic mice obtained by crossing Gt(ROSA)26Sor<tm14(CAG–tdTomato)Hze> reporters (Madisen et al., 2010) with appropriate drivers: *Pvalb*<tm1(cre)Arbr> (Hippenmeyer et al., 2005) (2 males, 3 females), *Vip*<tm1(cre)Zjh> (Taniguchi et al., 2011) (3 males, 2 females), *Sst*<tm2.1(cre)Zjh> (Taniguchi et al., 2011) (2 males, 1 female), and *Gad2*<tm2(cre)Zjh> (Taniguchi et al., 2011) (2 females). Experiments in which indicator was expressed uniquely in one neuron class were conducted in single transgenic mice: *Emx1*-IRES(cre) (n=1), *Pvalb*<tm1(cre)Zjh> (n=1), *Vip*<tm1(cre)Zjh> (n=1), *Sst*<tm2.1(cre)Zjh> (n=3), referred to as *Vip-Cre* and Sst-Cre respectively. Experiments in which pyramidal cells were labelled exclusively were conducted in *CamK2a–tTA;* Ai94(TITL– GCaMP6s); Rasgrf2–2A–dCre triple transgenic mice (n=3) (Madisen et al., 2015). Mice were used for experiments at adult postnatal ages (P54-110).

### Animal preparation and virus injection

The surgeries were performed in adult mice (P35-P76) in a stereotaxic frame and under isoflurane anesthesia (5% for induction, 0.5-1% during the surgery). During the surgery we implanted a head-plate for later head-fixation, made a craniotomy with a cranial window implant for optical access, and, on relevant experiments, performed virus injections, all during the same surgical procedure. In experiments where an interneuron class was recorded together with other cells, mice were injected with an unconditional GCaMP6m virus, AAV1.Syn.GCaMP6m.WPRE.SV40 (referred to as non-flex.GCaMP6m). In experiments where an interneuron class was labelled by unique expression, mice were injected with AAV1.Syn.Flex.GCaMP6m.WPRE.SV40 (flex.GCaMP6m) and AAV2/1.CAG.FLEX.tdTomato.WPRE.bGH (flex.tdTomato); all viruses were acquired from University of Pennsylvania Viral Vector Core. In both cases, viruses were injected with a beveled micropipette using a Nanoject II injector (Drummond Scientific Company, Broomall, PA 1) attached to a stereotaxic micromanipulator. One to three boli of 100-200 nl virus (2.23x10^12^ GC/ml for non-flex.GCaMP6m; 2.71x10^12^ for flex.GCaMP6m) were slowly (23 nl/min) injected unilaterally into monocular V1 (Wagor et al., 1980), 2.1-3.3 mm laterally and 3.5-4.0mm posteriorly from Bregma and at a depth of L2/3 (200-400 μm).

### Intrinsic Imaging

Prior to performing calcium imaging experiments, we performed intrinsic imaging of the optically accessible cortex to confirm the location of V1 within the cranial window (Supplementary Figure 1A,B). The intrinsic imaging was performed in all mice (n = 22) about 7-14 days after the surgery. We illuminated the cortex through the epi-illumination path using a high-power LED (central wavelength: 560 nm, M565L3, Thorlabs, Ely, UK), and acquired images at 5 Hz at 1024 x 1024 pixels using a CMOS camera (MV-D1024E-160; Photon-focus, Lachen, Switzerland) combined with a microscope objective (4x, NA: 0.13, FN: 26.5, UPLFLN, Olympus, Tokyo, Japan). To prevent the light contamination from the computer monitors we optically shielded the recording chamber with a custom black cone surrounding the objective.

### Retinotopic mapping from intrinsic imaging

To obtain retinotopic maps from intrinsic imaging we used the methods described in Pisauro et al. (2013). Briefly, we first removed global fluctuations from the signal, which are not stimulus driven. The residual signal reflects the retinotopic, stimulus-evoked responses. Visual stimuli were periodic drifting and flickering bars (Kalatsky and Stryker, 2003). Flickering bars (flicker frequency 2 Hz) drifted (speed = 0.8 deg/s) across -135° to 45° of the horizontal visual field (with bars oriented vertically) and -45° to 45°of the vertical visual field (with bars oriented horizontally) for 3 cycles. We calculated retinotopic maps using the method described in Kalatsky and Stryker (2003). Retinotopic contours (Supplementary Figure 1B) where obtained after removal of artefactual extreme values (e.g. red regions in the top and bottom left corners of Supplementary Figure 1A) and replacing the removed values by values interpolated using sum of normalized Gaussian functions with standard deviation of 20 m centered on non-artefactual pixels. Consistent with a location in V1, the imaged regions were within an area of diameter at least 2 mm where the gradient of vertical retinotopy was aligned from anterior to posterior (lower to higher values of elevation) and the gradient of the horizontal retinotopy was aligned from medial to lateral (temporal to central).

### Visual stimuli

Stimuli were horizontal gratings drifting downward, presented in a location adjusted to match the center of GCaMP expression, on one of two screens that together spanned -45° to +135° of the horizontal visual field and ±42.5° of the vertical visual field (left and central screens in Figure 1A_1_). During gray screen presentation, the screens were set to a steady gray level equal to the background of all the stimuli presented for visual responses protocols. Gratings had a duration of 1-2 s temporal frequency of 2 Hz and spatial frequency of 0.15 cycles/deg.

### Eye-tracking movie acquisition and analysis

For eye tracking we used a collimated infrared LED (SLS-0208-B, λ_peak_ = 850nm; controller: SLC-AA02-US; Mightex Systems, Toronto, Canada) to illuminate the eye contralateral to the recording site. Videos of eye position were captured at 30 Hz with a monochromatic camera (DMK 21BU04.H, The Imaging Source, Bremen, Germany) equipped with a zoom lens (MVL7000; Navitar, Rochester, NY), and positioned at approximately 50° azimuth and 50° elevation relative to the center of the mouse’ field of view. Contamination light from the monitors and the imaging laser was rejected using an optical band-pass filter (700-900nm) positioned in front of the camera objective (long-pass 092/52x0.75, The Imaging Source, Bremen, Germany; short-pass FES0900, Thorlabs, Ely UK).

To calibrate pupil displacement relative to the mouse visual field, we recorded additional movies at the end of each experiment while the mouse was still in exactly the same position as during the experiment. The eye was illuminated sequentially from a grid of known locations, the reflections were captured by the camera, and then this reflected grid was used to map the pupil displacement in pixels to pupil displacement in degrees of visual field.

Movie processing was performed offline using custom code written in MATLAB (Mathworks, Natick, MA) on a frame-by-frame basis. Briefly, each frame was mildly spatially low-pass filtered to reduce noise, then the pupil contour was detected by a level-crossing edge detector, and finally the position and the area of the pupil were calculated from the ellipse fit to the pupil contour. The output of the algorithm was visually inspected, and adjustments to the parameters (e.g. spatial filter strength, or level-crossing threshold) were made if necessary.

### In vivo calcium imaging

Experiments were performed 16-34 days after virus injection (P54-110). We used a commercial two-photon microscope (B-scope, ThorLabs, Ely UK), with an acquisition frame rate of about 30Hz (at 512 by 512 pixels, corresponding to a rate of 4-6 Hz per plane), which was later interpolated to a frequency of 10 Hz, common to all planes. Recordings were performed in the area where expression was strongest. In most recordings (n = 16) this location was in the monocular zone (MZ, horizontal visual field preference > 30°)(Wagor et al., 1980). Other recordings (n = 11) were performed in the callosal binocular zone (CBZ, n= 4, 0-15°)(Wang and Burkhalter, 2007) and others (n = 7) in the acallosal binocular zone (ABZ, 15-30°). We observed no difference in results between recordings in monocular and binocular zones (Supplementary Figure 12).

### Calcium data processing

Raw calcium movies were analyzed with Suite2p, which performs several processing stages (Pachitariu, 2016). First, Suite2p registers the movies to account for brain motion, then clusters neighboring pixels with similar timecourses into regions of interest (ROIs). ROIs were manually curated in the Suite2p GUI, to distinguish somata from dendritic processes based on their morphology. Cells expressing tdTomato were identified semi-automatically using an algorithm based on their average fluorescence in the red channel. For spike deconvolution from the Calcium traces, we used the default method in Suite2p (Pachitariu, 2016). Whether we performed performing spike deconvolution or analysed raw calcium signals made no difference to our results (Supplementary Figure 13).

### Pixel maps of calcium data

To confirm the correlation of running speed and fluorescence independent of ROI detection, we computed correlation maps (Supplementary Figure 2C), showing for each pixel the Pearson correlation between the activity of the pixel and the running speed (c.f. Freeman et al. (2014)). Prior to correlation, the activity of each pixel was smoothed by convolving with a spatial Gaussian with standard deviation equal to 1.5 pixels, and a temporal Hamming window of 1 s width.

The correlation of baseline fluorescence with running speed varied across the field of view. In regions where GCaMP expression level was high, baseline fluorescence correlated positively with running speed, likely indicating an increase in axonal and dendritic activity in locomoting animals. However, in areas where GCaMP fluorescence was weak, the correlation of the background with running speed was negative, likely indicating that in absence of GCaMP the signal is dominated by increased hemodynamic filtering of the light due to stronger blood flow during running (Huo et al., 2015). To ensure this did not affect our results, we removed background fluorescence from the detected fluorescence of recorded neurons (see below).

### Background fluorescence correction

With two-photon GCaMP imaging, an important concern is that out-of-focus fluorescence can contaminate the signal ascribed to particular neurons; this is of particular concern in situations where the surrounding GCaMP-labelled neuropil may itself show modulation by stimuli or behaviors such as locomotion. In order to correct out-of-focus contamination, we adopted the method of Peron et al. (2015). A “neuropil mask” was defined as the region up to 35 |im from the ROI border, excluding pixels corresponding to other detected cells (Supplementary Figure 2Ai), and the fluorescence signal in this mask region was subtracted from that of the cell soma, weighted by a correction factor *α_exp_* that was determined separately for each experiment.

To determine the correction factor, we estimated the linear relationship specifying the lowest possible somatic fluorescence compatible with any value of fluorescence in the neuropil mask (Supplementary Figure 2A_2_). To do so, for each cell *i* we binned the neuropil signals *N_i_(t)* into 20 intervals, and for each one estimated the 5^th^ percentile of the raw somatic fluorescence *F_i_(t)*. We computed *α_t_* by linear regression, which accurately matched the lower envelope of the scatterplot of neuropil vs. somatic fluorescence (Supplementary Figure 2A_2_). This method gave consistent results for sparse firing cells, but not always for densely firing cells for which a correlation of cellular activity with the neuropil signal could lead to misestimated slopes, as densely firing cells might only rarely exhibit baseline fluorescence. We therefore computed the correction factor *a_exp_* for each experiment by averaging *a_t_* over cells with high skewness ( > 4). The corrected fluorescence was computed as *F(t)* – *α_exp_N(t)* (Supplementary Figure 2A3). In experiments where only interneurons (and thus low skewed cells) expressed GCaMP6, we used as a correction factor an average from all the other experiments equal to 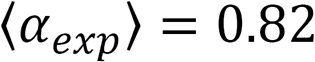.

### Analysis of neural activity

The average fluorescence response to each stimulus was defined by ΔF/F_0_ = (F-F_0_)/F_0_, where F is the average raw calcium signal during the first second of the stimulus presentation, and F0 is the global minimum of the fluorescence trace filtered with a Hamming window of duration 0.5 s. The correlation of neural activity with locomotion speed during gray screen presentation was assessed by the Pearson correlation coefficient between the calcium signal and the locomotion speed trace, on an interpolated timebase of 10 Hz, smoothed (5 points) and decimated (1 Hz). To ascertain the significance of this correlation we used a shuffling method, in which the speed trace was randomly circularly shifted relative to the fluorescence trace 1,000 times – this was necessary because serial correlation in the time series of fluorescence and speed rendered successive samples statistically dependent.

The size of a cell’s response to a stimulus was defined by the difference of ΔF/F_0_ between the first 1 s of the stimulus period, and the 1 s of baseline activity prior to stimulus presentation. We defined a neuron to have significant size tuning if it passed in at least one of the two locomotion conditions (rest or running) a one-way ANOVA test (p < 0.05) comparing the mean visual responses to different stimuli.

To measure each cell’s retinotopic location, in the majority of datasets (n = 24) receptive fields were obtained from responses to sparse, uncorrelated noise. The screen was divided into squares of 5 by 5 degrees, and each square was independently turned on and off randomly at a 5Hz overall rate. At any time, 5% of all squares were on. Each cell’s response to each square was obtained using stimulus-triggered averaging of the non-neuropil corrected trace. The RFs were smoothed in space and their peak was identified as the preferred spatial position. In a subset of early experiments (n = 3), sparse noise was not presented, and RFs were assessed with flickering vertical or horizontal bars appearing in different locations; we verified in a further n = 4 datasets that the two measures of a cell’s receptive field were consistent.

When computing size tuning curves, we normalized the calcium activity (Figure 3, column 3 and Supplementary Figure 12) or the spike rate (Supplementary Figure 13) in the following way: for each cell, the response Δ*F*_0_ to the “blank condition” (i.e. a stimulus of contrast 0) during stationary periods was subtracted from the raw cell response ΔF (1 s during stimulation minus 1 s of baseline activity): Δ*F* → Δ*F* − Δ*F*_0_. Then, for each recording we computed the average 〈Δ*F*〉_all cells_ over all selected cells whose distance of the receptive field from the stimulus center *r* was < 20°. Finally we divided the average response of either the centered cells (r < 10°) or the off-centered cells (r > 10°, r < 20°) by the maximum value of all the cells combined: 〈Δ*F*_*cent*._/max(〈ΔF〉_*all cells*_) or 〈Δ*F*_off–*cent*._/max(〈Δ*F*〉_*all cells*_) We then averaged these values across all recording sessions.

When computing the interaction between locomotion and size (Figure 3, column 4 and Supplementary Figure 10A,B) we normalized the responses of the individual cells for each experiment in the following way: after subtracting the average response to the blank during stationary periods Δ*F* Δ*F* – Δ*F*_0_, we divided the responses by the average minimal calcium trace across cells (*F*_0_)*_cells_* within the same experiment or mouse (Supplementary Figure 10A,B).

### Curve fitting

We fitted the size tuning curves of Figure 3 and Supplementary Figures 4, 8, 9, 12, and 13 by least squares with the following function family: *f(s)* = *R*[erf(*s*/(σ_1_) – *k* ⋅ erf(*s*/σ_2_)] where erf(*x*) corresponds to the error function and s is the size of the stimulus. The free parameters of the function are *R*, *k*, σ_1_ and *σ*_2_. To estimate nonlinear signal correlation curves (Figure 4, columns 3 and 4), we first smoothed the responses to large sizes (s >20°) for each population with a moving average method with span 25°. Then we smoothed again the responses for all sizes with a moving average method with span 20°. Finally we interpolated the values between the measured size with a shape-preserving piecewise cubic interpolation.

### Size tuning maps

To compute how size tuning depends on stimulus centering, we computed two-dimensional maps illustrating how each cell class’ average activity depends on stimulus diameter *s,* and the offset of the receptive field center from the stimulus center *r_i_* (Supplementary Figure 11). To do so, we first computed for each cell i a normalized tuning curve *n_iv_(s*), where *v* represents locomotion condition. Dependence on *r_t_* was estimated by smoothing: a two-dimensional map was made for each cell as an outer product: *f_i,v_(s,r)* = *n_i,v_*(*S*)*g_i_*(*r*), where 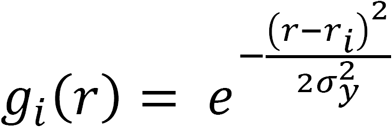 is a Gaussian centered at the offset value *r_i_* of width *σ_y_* = 5°. Then we summed the maps belonging to one recording session j and divided by the sum of all the Gaussians centered at different offsets: 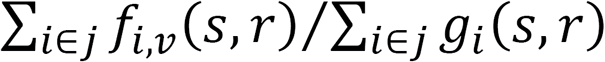, and normalized this value for each experiment by the value at stationary, 0 offset and diameter 10° *f_j,v_(s,r)* = *m_j,v_(s,r)/m_j,v=1_(s* = 10,*r* = 0). Finally, for each cell class we obtained the size-tuning offset maps by averaging across experiments: 〈*f_j,v_*(*s,r*))〉.

### Inter-population correlation analysis

To compute spontaneous correlations (Figure 4, columns 1 and 2), we first normalized the deconvolved spike trace *S* of each cell *i* over time *t: n(i,t) = S(i, t)/max_t_(S).* Then, for each experiment and each class *c* (interneurons or putative Pyr cells) we computed the average population rate across cells *i* belonging to class c: 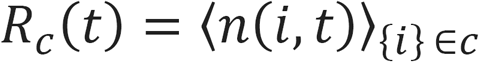. We then smoothed *R_c_(t)* with a moving average method with span of 1 s and then decimated the sampling rate to 1 point every 1 s. To make the plots of different experiments visually comparable we normalized these responses: 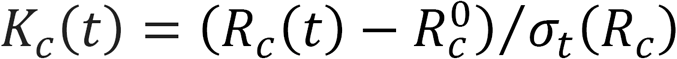 Where is the 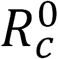 percentile of *R_c_* and *σ_t_(R_c_)* the standard deviation of *R_c_(t)* across time.

To compute signal correlations (Figure 4, columns 3 and 4), for each experiment and each cell class *c* we first computed the average population response for each stimulus size *s* and locomotion condition *v* (*v* = 0 stationary, *v* = 1 running) by averaging over all cells *i* belonging to that class: 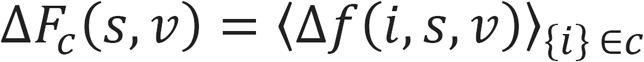. To make the plots of different experiments visually comparable we normalized the responses: Δ*R_c_(s,v)* = Δ*F_c_(s,v)* /σ_s,v_(Δ*F_c_*). Finally we subtracted the blank response during the resting condition: Δ*K_c_(s,v)* = Δ*R_c_(s, v)* — Δ*R_c_*(*s* = 0,*v* = 0).

To compute noise correlations (Supplementary Figure 14, columns 1 and 2), for each experiment and each cell class *c* we first computed the average population response for each trial *t* by averaging over all cells *i* belonging to that class: 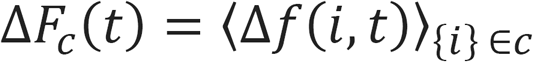. Then for each stimulus and locomotion condition we subtracted the mean response from the related trials: 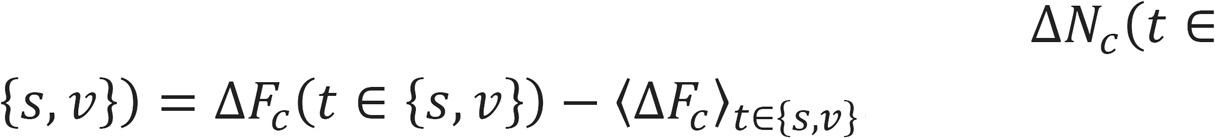 Finally, to make the plots of different experiments visually comparable, we normalized the responses by z-scoring over all trials: Δ*Z_c_(t)* = (Δ*N_c_(t)* — 〈Δ*N_c_*〉_t_)/σ_t_(Δ*N_c_*).

When measuring signal and noise correlations and for both interneurons and putative Pyr neurons, we selected cells whose receptive field center was <10° away from the stimulus center. For Figure 4 we selected putative Pyr cells as unlabeled (non tdTomato) neurons whose skewness was > 2.7. In a control analysis (Supplementary Figure 15) we show that the value of the skewness threshold makes little difference to these results. A skewness value of 0 corresponds to the case where we selected all unlabeled cells as putative Pyr cells.

### Correlation of running modulation with depth

To determine whether running modulation of a given cell class varied significantly with cortical depth, we computed *ρ*(*gray*) for each cell as the Pearson correlation of that cell’s neuropil-corrected fluorescence (without spike deconvolution) and running speed. We then assessed a significant relationship of *ρ*(*gray*) with depth using robust regression (bisquare-weighting).

### Computational model

We asked whether we could predict the mean size tuning of each cell class using a neural field theory model. In this model, the mean firing rate of cells of type *α* is captured by a function *f_α_*(*s*,**r**), where *s* represents the stimulus size, and **r** represents position on the cortical surface, measured in retinotopic coordinates. We model the external excitatory input arriving at point *r*
(e.g. from thalamus or other cortical layers) by a function *h(s,* **r**), again in retinotopic coordinates.

We denote the experimentally measured responses of cell class *α* by 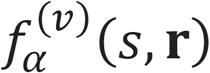, where *v* denotes locomotion condition (*v* = 0: stationary, *v* = 1: running). We assume that responses are circularly symmetrical, i.e. that responses depend on **r** only through the radial distance of the receptive field center from the stimulus center, *r*. The response of each cell class 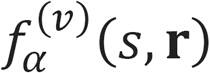 is modeled by the following equations:

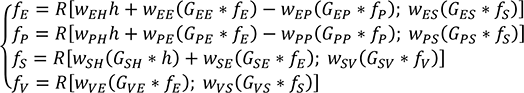

Only synaptic connections demonstrated in vitro (Pfeffer et al., 2013) are included in this equation; however, adding other potential synapses (Vip–>Pyr) did not improve the fit (Supplementary Figure 16).

For each postsynaptic cell class we tested different combination of subtractive and divisive inhibition from Sst and Vip cells:

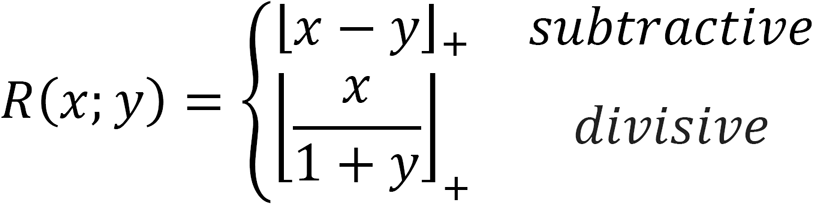

Here, *f_E_, f_P_ f_S_*, and *f_v_* reflect the visual responses of the Pyr, *Pvalb, Sst*, and *Vip* cells respectively; 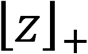 is the positive part of *z*; *w_αβ_* are the peak synaptic weights between the presynaptic cell class *β* and the postsynaptic cell class *α* (which can in principle depend on running condition *v*); *G_αβ_* is a two-dimensional Gaussian function defined by *G_αβ_*(Z) = exp[−|z|^2^/(2*σ^2^_αβ_*)]/(2*πσ^2^_αβ_*, with radius *σ_αβ_* that can depend on the pre- and post-sysnpatic cell type; and * represents convolution over retinotopic space: 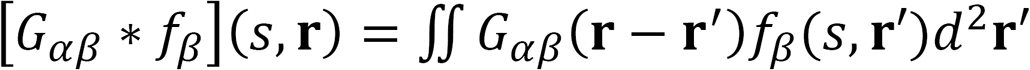

The equations describing the activity of the cell classes can be simplified with the following assumptions:

- 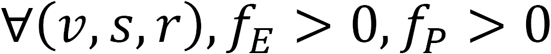(Pyr and *Pvalb* cells are not suppressed by stimuli, as seen in the data).
- The recurrent connections of Pyr and *Pvalb* neurons, and the connection from *Pvalb* to Pyr are local: *G_EE_(r) = G_EP_(r) = G_pp_(r) = δ(r)* where *δ(r)* is the Dirac delta function.
- 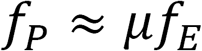 (i.e. Pvalb activity closely tracks Pyr activity, as seen in the data).

We can then rewrite the equations as:

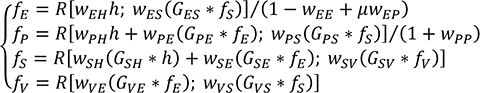

We can further simplify this equation as:

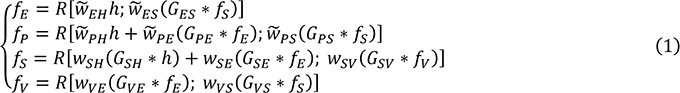

where 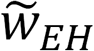 and 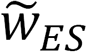 are the “effective connections” from external inputs and *Sst* neurons to excitatory cells, given by *W_EH_*/(1-*W_EE_*+*μW_EP_*) and *W_EH_*/(1-*W_EE_*+*μW_EP_*) represent the effective weights onto *Pvalb* cells, equal to *W_PH_*/(1+*W_pp_*), *W_PE_*/(1+*W_pp_*), and *W_PS_*/(1+*W_PP_*).

### Estimation of thalamic input

We estimated the average external inputs *h(s,* 0) from the experimentally-measured responses of centered (*r* = 0) thalamic cells using the data in Erisken et al. (2014), for stationary and locomotion periods separately. We first fit the size tuning curve of each cell (n = 21) with a ratio of Gaussians 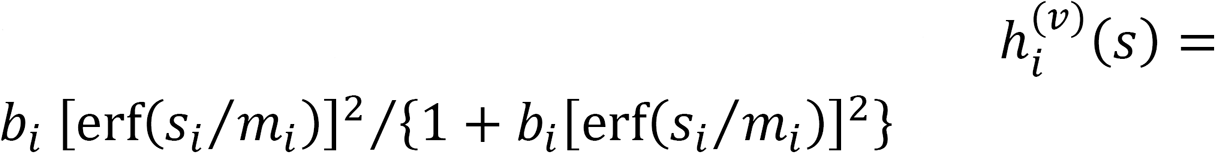 as in Erisken et al. (2014). We then normalized each 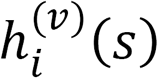 by its maximum across *s* and *v,* and estimated the mean visual responses by averaging across all cells. To fit off-center responses, we extended this approach using a Ratio-of-Gaussians model (Ayaz et al., 2013): *h(s,r)* = *a_1_ u(s,r,σ*_1_)/[1 + *a_2_u(s,r,σ_2_*)] with *u(s,r,σ)* = erf[(*s + r*)/σ] + sign(*s – r*)erf(|s – r|/σ). The estimated parameters are during the stationary periods were *a_1_* = 1.2, *a_2_* = 1.9, *r_1_* = 36.7, and *r_2_* = 33.9 while during locomotion *a_1_* = 0.5, *a_2_* = 0.4, *σ_1_* = 24.7°, and *σ_2_* = 10.0°.

### Estimation of presynaptic inputs

To fit the model, we clamped the firing rate functions 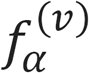 in equation (1) to their experimentally-measured values, and fit synaptic parameters to reduce the discrepancy between the right and left sides. To do so required extending our experimental data to continuous functions of s and *r.* For retinotopic positions *r* < *r_L_* = 33°, *f(s,r)* was fitted from the data with a difference of Gaussians function: *f(s,r)* = *R_r_*[erf(*s/σ*_1,r_) – *k_r_* • erf(*s/σ*_2,s_)]. Because our data for cells off-center by more than 33° were sparse, we extrapolated the values for *r* > *r_L_* with a decaying exponential approximation: 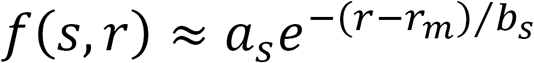 where *r_m_* is the offset value that maximizes the response of that cell class, and the parameters *a_s_* and *b_s_* were estimated for each stimulus size to minimize squared error in the range 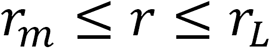 Finally, the values of *b_s_* were interpolated beyond the presented stimulus sizes with a third degree polynomial function.

### Parameter estimation

To estimate the parameters we minimized an objective function equal to the normalized mean-square error, plus additional penalty terms to favor simpler models:

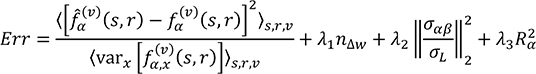

Here, 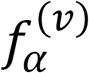 represents the measured firing rate, and 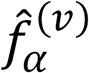 represents the right hand size of equation (1). The normalization factor of 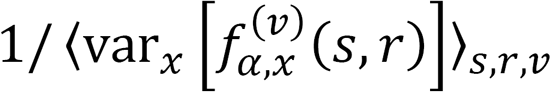 ensures that conditions with high intertrial variability do not overly influence the objective function; the normalized error can also be interpreted as the log-likelihood of the model fit under a Gaussian distribution estimated from all experiments 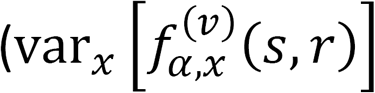 denotes the variance of normalized visual responses for each value of *s*, *r* and *v* over all experiments *x*).

The last three terms represent regularization parameters. The first regularization term controls the number of synaptic strengths that are allowed to change with locomotion, *n_Δw_*; for the current analysis we used a value of *λ*_1_ = 0.1. The second regularization term controls the spatial distribution of synaptic weights; we used parameters *σ_L_* = 40°, *λ*_2_ = 0.01. The final term *R_α_*, with *λ*_3_ = 0.2, represents a factor to add biologically motivated constraints in the *Pvalb* and *Sst* equations.

We assume that Sst cells receive most of their input from a local area, therefore 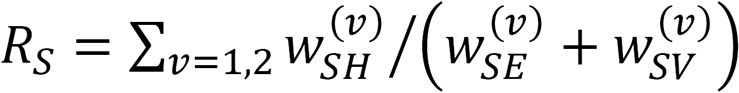.

The activities of Pyr and Pvalb are very similar, so to avoid *f_E_* dominating the *f_P_* equation we have 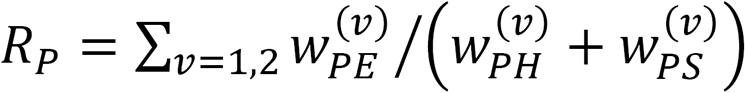.

To determine the optimal parameters of the model we first performed an exhaustive search over the extent of the spatial integration *σ_βα_* parameters of all the presynaptic cell classes for each postsynaptic cell class. For each combination of 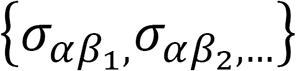 we then found the optimal synaptic strength parameters 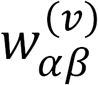 using a combination of the trust region reflective and Levenberg-Marquardt algorithms (MATLAB), after 50 random initialization of the initial parameters 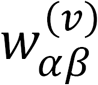. We then chose the values of *σ_αβ_* and 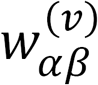 minimizing *E_rr_*.

To fit the way locomotion affects synaptic strengths, we found the optimal of several possibilities for each synaptic type: 1) equal weights in locomotion and stationary 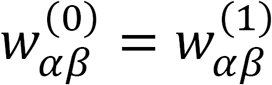; 2) fixing all but one of the synaptic weights 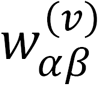 (therefore testing one model for each synaptic connection) 3) allowing all the synaptic weights to change with locomotion. The final model was chosen as that minimizing the penalized error function *Err* (Supplementary Figure 16A-D1).

## Author contributions

M.D. performed the experiments, analyzed the data and implemented the model; M.D., A.R. and M.P. performed the surgeries; A.R. and M.K. built the two-photon set-up; M.D, M.K. and M.P. implemented the retinotopic mapping software; K.D.H. and M.P. implemented the ROI detection algorithms; M.D., A.R., M.C. and K.D.H. designed the experiments; M.D. and K.D.H. designed the model; M.D., M.C. and K.D.H. wrote the paper with inputs from all authors.

## Acknowledgments

We thank Charu Reddy for transgenic mouse breeding and maintenance, surgeries, and technical support, Chris Burgess for building the data acquisition software set-up, Luigi Federico Rossi for assistance during surgeries, Miles Wells for assistance in mice habituation,Andrea Pisauro for implementing the intrinsic imaginc retinotopic mapping software, Sylvia Schroeder for implementing the calcium retinotopic mapping software. We thank Boris S. Gutkin for constructive discussion. For the recorded size tuning curves of the dLGN cells in vivo we acknowledge Laura Busse’s lab. For the use of GCaMP6f we acknowledge Vivek Jayaraman, Rex A. Kerr, Douglas S. Kim, Loren L. Looger, Karel Svoboda from the GENIE Project, Janelia Farm Research Campus, Howard Hughes Medical Institute. This work was supported by the Wellcome Trust. MD was supported by a Marie Curie Intra-European Fellowship for Career Development, KDH by EPSRC. MC holds the GlaxoSmithKline / Fight for Sight Chair in Visual Neuroscience.

